# Sequence-based Protein-Protein Interaction Prediction Using Multi-kernel Deep Convolutional Neural Networks with Protein Language Model

**DOI:** 10.1101/2023.10.03.560728

**Authors:** Thanh Hai Dang, Tien Anh Vu

## Abstract

Predicting protein-protein interactions (PPIs) using only sequence information represents a fundamental problem in biology. In the past five years, a wide range of state-of-the-art deep learning models have been developed to address the computational prediction of PPIs based on sequences. Convolutional neural networks (CNNs) are widely adopted in these model architectures; however, the design of a deep and wide CNN architecture that comprehensively extracts interaction features from pairs of proteins is not well studied. Despite the development of several protein language models that distill the knowledge of evolutionary, structural, and functional information from gigantic protein sequence databases, no studies have integrated the amino acid embeddings of the protein language model for encoding protein sequences.In this study, we introduces a novel hybrid classifier, xCAPT5, which combines the deep multi-kernel convolutional accumulated pooling siamese neural network (CAPT5) and the XGBoost model (x) to enhance interaction prediction. The CAPT5 utilizes multi-deep convolutional channels with varying kernel sizes in the Siamese architecture, enabling the capture of small- and large-scale local features. By concatenating max and average pooling features in a depth-wise manner, CAPT5 effectively learns crucial features with low computational cost. This study is the first to extract information-rich amino acid embedding from a protein language model by a deep convolutional network, through training to obtain discriminant representations of protein sequence pairs that are fed into XGBoost for predicting PPIs. Experimental results demonstrate that xCAPT5 outperforms several state-of-the-art methods on binary PPI prediction, including generalized PPI on intra-species, cross-species, inter-species, and stringent similarity tasks. The implementation of our framework is available at https://github.com/anhvt00/MCAPS

## Introduction

In the complex cellular environment, proteins regularly interact with each other, providing the foundation for numerous vital biological functions. These interactions, called protein-protein interactions (PPIs), act as regulatory hubs for a diverse range of cellular processes, including gene expression, cell signaling, and metabolic pathways. To recognize and analyze PPIs, a wide variety of experimental methods, both high-throughput and low-throughput, have been created. Nevertheless, these techniques are often hindered by their high cost, time-intensive nature, and limited accuracy. The field of computational biology has witnessed the emergence of various methods to predict PPIs. These computational approaches have the potential to infer a large number of PPIs with a high degree of accuracy. A substantial portion of these methods is focused on predicting PPIs solely through protein sequences.

Deep learning models have emerged as the vanguard of computational approaches for binary PPI prediction. Notable among these is DPPI (1), which employs a deep Siamese-like convolutional neural network with random projection and data augmentation, using PSI-BLAST (2) protein representations as inputs. DPPI has the distinction of being the first deep learning model to achieve state-of-the-art performance in binary PPI prediction. PIPR (3) employs a Siamese architecture and a residual recurrent convolutional neural network (RCNN) to capture both local significant features and sequential features, providing an automatic multi-granular feature selection mechanism.

D-SCRIPT (4) is a deep-learning model that predicts protein-protein interactions directly from protein sequences. It has two stages: the first stage generates rich features for each protein, and the second stage predicts an interaction based on these features. D-SCRIPT innovative is in its structurally aware design, which encodes a physical model of protein interaction to predict how proteins interact and form pathways. FSNN-LGBM (5) is a hybrid classifier that combines a functional-link-based neural network (FSNN) and a LightGBM boosting classifier. DeepTrio (6) follows a mask multiscale CNN architecture that captures multiscale contextual protein sequence information using multiple parallel filters. Topsy-Turvy (7) is a computational method that combines both sequence-based and global network-based views of protein interaction. During training, it takes a transfer-learning approach by incorporating patterns from both global and molecular-level views, resulting in state-of-the-art performance in PPI prediction.

TAGPPI (8) incorporates both sequence features and predicted structural information and employs graph representation learning methods on contact maps to obtain 3D structure features of proteins. HNSPPI (9) adopts a feature fusion strategy of both network topology and sequence information for comprehensive feature extraction and employs a simple classifier for predictions, making it both lightweight and efficient. Graph-BERT (10) utilizes a language model-based embedding SeqVec to represent protein sequences and a graph convolutional neural network with the training strategy of subgraph batches using a top-k intimacy sampling approach. The MARPPI model (11) is a multi-scale architecture residual network designed for predicting Protein-Protein Interactions (PPIs). It employs a dual-channel and multi-feature approach, leveraging Res2vec for association information between residues and utilizing pseudo amino acid composition, autocorrelation descriptors, and multivariate mutual information for comprehensive feature extraction.

## Materials and Methods

### Model Architecture

In this section, we present the general architecture of our xCAPT5 model, which combines the neural network xCAPT5 with the boosted model XGBoost for sequence-based binary PPIs prediction. xCAPT5’s architecture is depicted in Figure 1. In general, our model encompasses five distinct phases, namely Amino Acids (AA) encoding layer, protein sequence learning layer, protein pair learning layer, intermediate layer, and prediction layer. Each layer plays a crucial role in the overall architecture and functionality of xCAPT5. xCAPT5 utilizes a deep multi-kernel convolutional neural network in a Siamese architecture to effectively capture and leverage the mutual latent representations between two protein sequences, which can be used to predict PPIs. Protein sequences are represented as amino acid embeddings using the Protein Language Model ProtT5XL-UniRef50 (12), which captures various protein properties including evolutionary, physicochemical, and structural information.

**Fig. 1.**
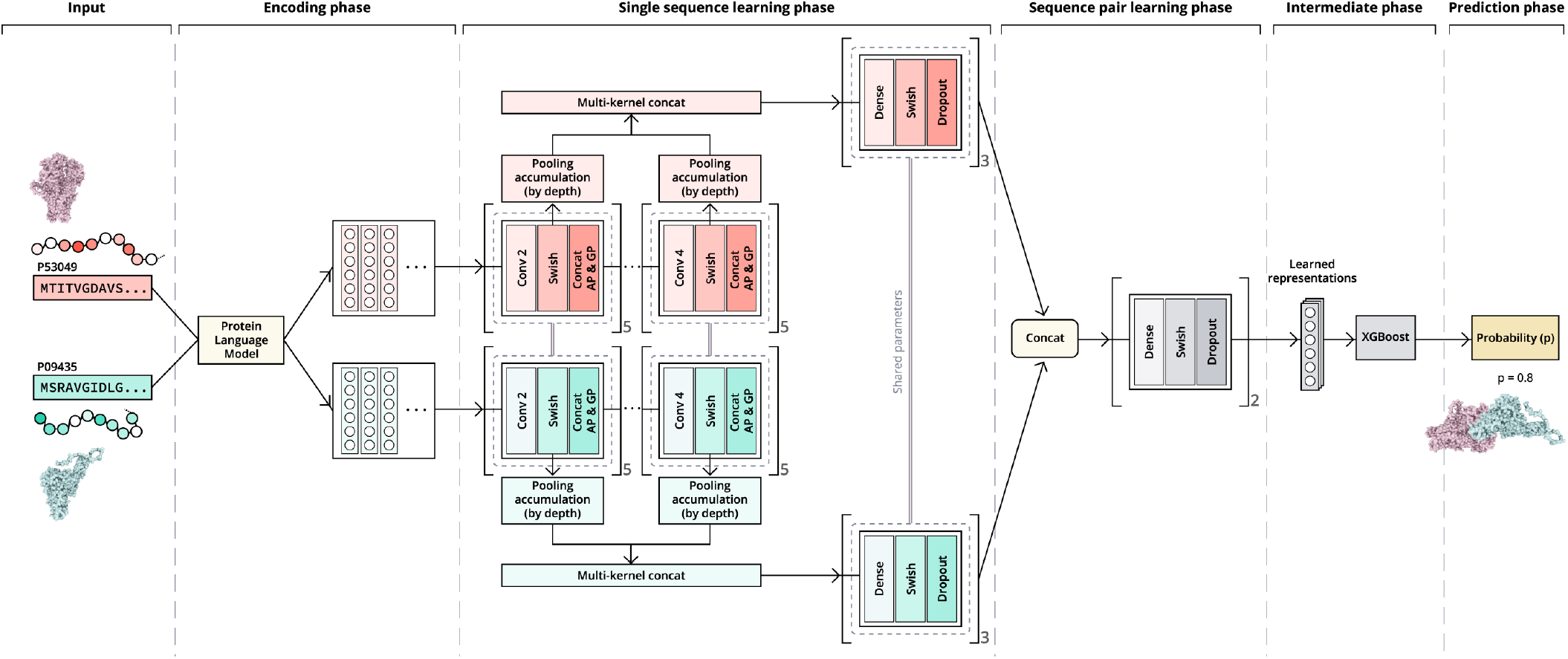
General architecture of xCAPT5 model.

#### Encoding Phase

To generate the amino acid embeddings, xCAPT5 employs the Protein Language Model ProtT5-XL-UniRef50. This model effectively captures the nuanced relationships and features within protein sequences. By leveraging its pre-trained knowledge and understanding of protein structures and functions, ProtT5-XL-UniRef50 maps each amino acid in the input sequences to a high-dimensional representation. xCAPT5 transforms a pair of protein sequences, denoted as *S* and *S*′, into corresponding amino acid embeddings represented by *X* and *X*′. These embeddings are structured as matrices, where *X* and *X*′ have dimensions of ℝ^1200×1024^, encode crucial information about the amino acid composition and order in the respective protein sequences.

#### Protein Sequence Learning Phase

Following the encoding phase, the protein sequence learning phase in xCAPT5 delves into extracting and comprehending the intricate patterns and representations inherent within pairs of amino acid embeddings, *X* and *X*′. To achieve this, xCAPT5 employs a Siamese architecture that utilizes deep multi-kernel convolutional neural networks (CNNs) combined with the concatenation of global average pooling (GAP) and global max pooling (GMP).

The Siamese architecture is employed to process two protein sequences simultaneously, capturing their respective patterns and representations in a shared network. This architecture facilitates the learning of the latent relationships and interactions between the individual protein sequences. Within the Siamese architecture, the deep multi-kernel CNNs serve as the backbone for extracting meaningful features from the protein sequences. These CNNs employ multiple convolutional kernels, each with a different size *k* ∈ [2, 3, 4], to capture both local and global features. The multi-kernel approach enables the network to explore and learn diverse spatial relationships and motifs within the protein sequences, enhancing its ability to comprehend the complex characteristics embedded within them. To extract and capture the intricate information embedded within the rich-information amino acid embeddings, xCAPT5 constructs deep CNNs corresponding to each kernel size. The deep CNN within xCAPT5 is structured with five sequential layers, each representing a level of depth in the network.

### The first layer (Convolutional Layer)

applies a set of filters with kernel size *k* to the input *X* (the amino acid embeddings) in the first module or the output of the third layer from the previous module d-th 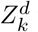 with *d* ∈ [1, 4], we denote 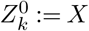. These filters capture different local patterns and interactions, allowing the network to detect important features within the protein sequences

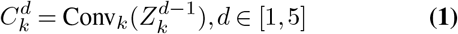

### The second layer (Swish Activation Layer)

introduces non-linearity into the network via the swish activation function (13). This function enables the model to capture intricate relationships and dependencies among the learned features effectively. The layer maps the feature maps 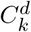 generated by the convolutional operations in the preceding layer to a set of activated feature maps 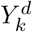.

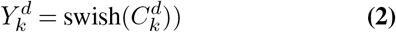

### The third layer (concatenation of average pooling (AP) and max pooling (MP))

receives the activated feature maps 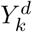 as inputs and performs both AP and MP operations followed by a spatial dropout operation, referred to as SpatialDrop, a regularization technique that randomly deactivates entire feature maps during training to prevents the model from relying excessively on specific spatial locations or local patterns, thereby reducing overfitting. This layer effectively combines global context information, derived from AP, and the most discriminative local features, derived from MP. Following the pooling operations, another spatial dropout operation is applied to further enhance the robustness of the model. The output of this layer is a set of pooled and regularized feature maps 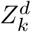.

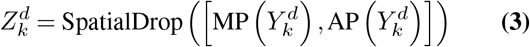

### The Fourth Layer (Pooling Accumulation)

not a direct layer in the flow of information through the deep CNN, instead it functions as a sidechain module. The GMP and GAP operations are applied to the output from the second layer 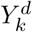, producing two vectors that represent the most significant (GMP) and average (GAP) features. These two vectors are then concatenated to form a comprehensive feature map that carries both global and local information about the input, which is then subjected to a dropout operation (denoted by Drop) to reduce overfitting.

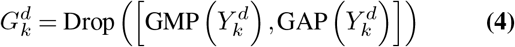

Consequently, the vectors 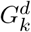 that are generated at each depth are accumulated in a depth-wise manner. This depth-wise accumulation ensures a comprehensive aggregation of information from all levels of the network. As a result, the module efficiently manages and integrates the critical feature information that has been extracted and processed by the previous layers in the deep CNN. This procedure facilitates a depth-wise understanding of the hierarchical representations of the protein sequences, thereby enhancing the model’s ability to interpret and learn from complex protein sequence data.

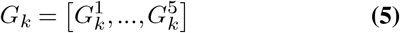

After the depth-wise pooling accumulation for each kernel size *k*, the resulting vectors *G*_*k*_ are concatenated. This comprehensive representation, denoted as *G*, captures a wide array of features from the input sequences. The vector *G* ∈ ℝ^1200^ is a fusion of information extracted by convolutional layers with different kernel sizes. We apply the batch normalization (BatchNorm) and the dropout operation follow to make the training more stable and generalize better.

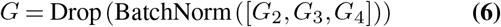

Deep CNN with different multiple kernel size working together allows the model to capture different scales of spatial relationships in the input data. Smaller kernel sizes can capture fine-grained, local features, while larger kernel sizes can pick up on more global, abstract features. By concatenating the accumulated vectors for each kernel size, the model can retain and leverage these diverse scales of features simultaneously. Upon capturing the features from the protein sequences through the convolutional neural networks (CNNs), these features embodied in the vector *G* are directed into a feed-forward block for further refinement and transformation. This process entails the application of linear transformations along with non-linear activation functions within the feed-forward block. As a result, the model is capable of encapsulating the vital characteristics of the protein sequence more effectively, contributing to a reduction in data dimensionality. The Siamese architecture ensures that both sequences in the pair go through the same processing steps with shared weights. This means that for the second sequence in the pair, a feature tensor *G*′ is created in the same way as *G* was for the first sequence. Both sequences are independently fed through the same deep multi-kernel CNNs, and the extracted features from each are then passed through the same feed-forward block. For each sequence, the output from the feed-forward block is a vector *S* ∈ ℝ^186^ or *S*′ ∈ ℝ^186^, depending on whether it’s the first or second sequence in the pair. The feed-forward block comprises three consecutive layers, each with a fully connected layer followed by a swish activation function and dropout. Here, *W*_1_ ∈ ℝ^744×1200^, *b*_1_ ∈ ℝ^744^, *W*_2_ ∈ℝ^372×744^, *b*_2_ ∈ ℝ^372^, *W*_3_ ∈ ℝ^186×372^, *b*_3_ ∈ℝ^186^ denote the weights and biases of the first, second, and third layer, respectively.

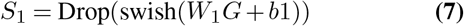

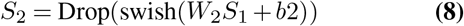

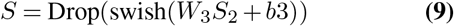

#### Sequence Pair Learning Phase

In the sequence pair learning phase, the goal is to capture the dependencies and characteristics that define the interaction between two protein sequences. To achieve this, the processed features of the two sequences, denoted as *S* and *S*^*/*^, are combined and fed into a multi-layer perceptron (MLP). This phase is crucial for learning the latent relationships and interactions between the pair, enabling accurate prediction of their interaction. To form a composite feature map, the refined feature vectors *S* and *S*′ are concatenated, resulting in a combined feature map 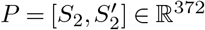. This composite feature map captures the information from both sequences and their potential mutual information. This concatenated feature map is then passed through a multi-layer perceptron (MLP), which composes of two densely connected layers, each followed by a swish activation function and a dropout operation. Here, *M*_1_ ∈ ℝ^328×372^, *c*_1_ ∈ ℝ^328^, *M*_2_ ∈ ℝ^164×328^, *c*_2_ ∈ ℝ^164^, *M*_3_ ∈ ℝ^1×164^, *c*_3_ ∈ ℝ denote the weights and biases of the first-, the second fully connected layer, and the output layer respectively.

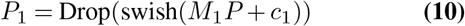

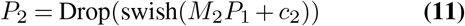

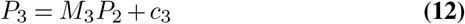

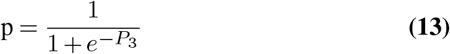

these equations illustrate the transformations that the combined feature map undergoes as it is passed through the MLP. The final output of the MLP, represented as p, is obtained by applying a sigmoid function to the output of the final dense layer. This sigmoid function maps the final output to a range between 0 and 1, thus making it interpretable as the probability of interaction between the protein sequence pair.

#### The intermediate phase

Subsequent to the initial training phase of the neural network xCAPT5, the derived representations from xCAPT5 are put into use. Once training is complete, the dataset is passed through xCAPT5 and the model’s penultimate layer representations, denoted as *P* , are extracted. These derived representations, *P* , are then fed into a XGBoost (14), a powerful gradient boosting framework, proceeds to further refine these representations, enhancing the model’s ability to capture complex patterns in the data. This additional layer of processing serves to enhance the model’s overall predictive power and accuracy.

#### Prediction Phase

Once the XGBoost model is fully trained, it can be used to predict PPIs. The model outputs a score for each protein pair, which can be interpreted as the predicted probability of interaction for that pair. A decision threshold is set, often at 0.5 for binary classification tasks. If the predicted probability is greater than this threshold, the model predicts that the pair of sequences interact. If the predicted probability is lower than the threshold, the model predicts that they do not interact. By leveraging the strengths of both deep learning through xCAPT5 and gradient boosting through XG-Boost, the model is able to effectively learn from the protein sequence data and accurately predict protein-protein interactions.

### Model Hyperparameters

We use three kernel size of 2, 3, 4. For each kernel size, each CNN is designed with a depth of 5 modules. The network employs a spatial dropout rate of 0.15 and a standard dropout rate of 0.05 to prevent overfitting and enhance generalization. We configure the hidden layers with 744, 372, and 186 units, while the final multilayer perceptron (MLP) after the merge has 328 and 164 units. For the optimization, we employ the Adam optimizer (15) with learning rate 1e-3, Amsgrad setting (**?** ), epsilon 1e-6, and batch size 64.

Regarding the XGBoost, the gbtree booster is used for utilizing gradient boosting trees. Regularization is applied via a reg_lambda (L2 regularization term on weights) of 1 and an alpha value (L1 regularization term on weights) of 1e-7 to prevent overfitting. Subsampling of the dataset and column sampling by tree are set at 0.8 and 0.2 respectively. The model utilizes 1000 estimators with a maximum tree depth of 5 to ensure a balance between model complexity and performance. The model also sets a minimum child weight of 2 to avoid overfitting. Furthermore, gamma of 1e-7 is used as a minimum loss reduction parameter and eta of 1e-6 as a learning rate to maintain a slow and steady model learning process.

### Datasets and Experiments

In this paper, we did three intensively thorough experiments to evaluate the performance of our model, comparing it with recent state-of-the-art PPI prediction models on several benchmark datasets. The evaluation metrics used were accuracy, precision, recall, specificity, F1-score, and Matthews correlation coefficient (MCC), Area Under the Receiver Operating Characteristic curve (AUROC), and Area Under the Precision-Recall curve (AUPRC).

The first experiment involves evaluating the learning capacity of models by conducting five-fold cross-validation on three golden standard datasets. These datasets include the Martin *H. pylori* dataset (16) with 1458 positive pairs and 1365 negative pairs, the Guo yeast dataset (17) with 5594 positive pairs and 5594 negative pairs, and the Pan human dataset (18) with 27593 positive pairs and 34298 negative pairs.

The second experiment focuses on evaluating the generalized inference capacity of models on three tasks: intra-species inference, cross-species inference, and inter-species inference. For intra-species evaluation, we use three human PPI datasets from Li’s work (19): HPRD with 3516 PPIs, DIP with 1468 PPIs, and HIPPIE HQ (high-quality) with 15489 PPIs, and HIPPIE LQ (low-quality) with 101684 PPIs. Cross-species evaluation involves testing the models on datasets from other species, including mouse, fly, yeast, *C. elegans*, and *E. coli*, retrieved from Sledzieski’s datasets (4). These datasets consist of 5000 positive pairs and 50000 negative pairs, except for the *E. coli* dataset, which has 2000 positive pairs and 20000 negative pairs. The inter-species evaluation focuses on human-other species PPI test datasets from Yang’s work (20). The datasets involve 8 viruses: HIV with 9880 positive pairs and 98800 negative pairs, Herpes with 5966 positive pairs and 59660 negative pairs, Papilloma with 5099 positive pairs and 50990 negative pairs, Influenza with 3044 positive pairs and 30440 negative pairs, Hepatitis with 1300 positive pairs and 13000 negative pairs, Dengue with 927 positive pairs and 9270 negative pairs, Zika with 709 positive pairs and 7090 negative pairs, and Sars-CoV-2 with 586 positive pairs and 5860 negative pairs.

The third experiment involves evaluating the learning capacity of models on more constrained datasets with different stringent similarity in sequences. Chen’s multispecies dataset ((3)) is used, with stringent similarity values ranging from 0.01 to 0.4. The performance of the models is evaluated using five-fold cross-validation, with higher stringent similarity values indicating more challenging datasets.

Our proposed xCAPT5 model is compared with five state-of-the-art models, including PIPR, FSNN-LGBM, D-SCRIPT, Topsy-Turvy, and DeepTrio. We note that Topsy-Turvy and D-SCRIPT are only compared in the second experiment due to computational constraints.

## Results

### Cross-validation performance

On the Martin data set (Table 1), xCAPT5 exhibits a consistent superior performance across various performance metrics. The model leads with an outstanding accuracy of 97.27%, significantly 1% higher than its closest competitor, FSNN-LGBM of 96.49%. xCAPT5 also excels in other metrics such as precision of 97.30%, specificity of 97.44%, F1-Score of 97.19, and Matthews Correlation Coefficient (MCC) of 94.82%. Interestingly, while HNSPPI shows a marginally better recall score of 99.39%, it falls short in other metrics like precision and MCC. This suggests that while HNSPPI is excellent at identifying true positives, it may not be as well-rounded as xCAPT5, which exhibits high performance in multiple metrics simultaneously.

**Table 1.** 5-Fold cross-validation performances of methods on Martin dataset.

| Method | Accuracy (%) | Precision (%) | Recall (%) | Specificity (%) | F1-Score (%) | MCC (%) |
| --- | --- | --- | --- | --- | --- | --- |
| PIPR | 80.84 ± 0.44 | 81.44 ± 0.69 | 81.55 ± 0.85 | 80.32 ± 0.67 | 81.43 ± 0.45 | 61.69 ± 0.89 |
| FSNN-LGBM | 96.49 ± 0.13 | 96.03 ± 0.26 | 97.23 ± 0.04 | 95.69 ± 0.29 | 96.62 ± 0.12 | 92.98 ± 0.25 |
| MARPPI | 91.80 ± 1.16 | 90.69 ± 2.68 | 94.51 ± 1.13 | 91.22 ± 1.25 | NA | 83.74 ± 2.32 |
| HNSPPI | 93.21 ± 0.35 | 88.47 ± 0.53 | **99.39 ± 0.21** | NA | 93.59 ± 0.32 | 93.21 ± 0.35 |
| Our xCAPT5 | **97.27 ± 0.12** | **97.30 ± 0.24** | 97.07 ± 0.20 | **97.44 ± 0.11** | **97.18 ± 0.25** | **94.82 ± 0.20** |

Report with mean and sd

Experimental results on the Guo data set demonstrate that xCAPT5 outperforms all compared models by significant margins across multiple key metrics. With a remarkable accuracy of 99.76%, xCAPT5 eclipses its nearest competitor, HNSPPI, which scored 98.57% in accuracy. (Table 2).In terms of precision, xCAPT5 maintains its dominion with a score of 99.76%, compared to FSNN-LGBM’s 98.73%, once again indicating superior specificity. The model’s recall rate is 99.75%, making it the leader in identifying true positive cases as well; the closest competitor here is HNSPPI at 98.85%. The same trend is evident in the specificity, F1-score, and Matthews Correlation Coefficient (MCC) categories, where xCAPT5 posts scores of 99.77%, 99.37% and 99.52%, respectively.

**Table 2.**
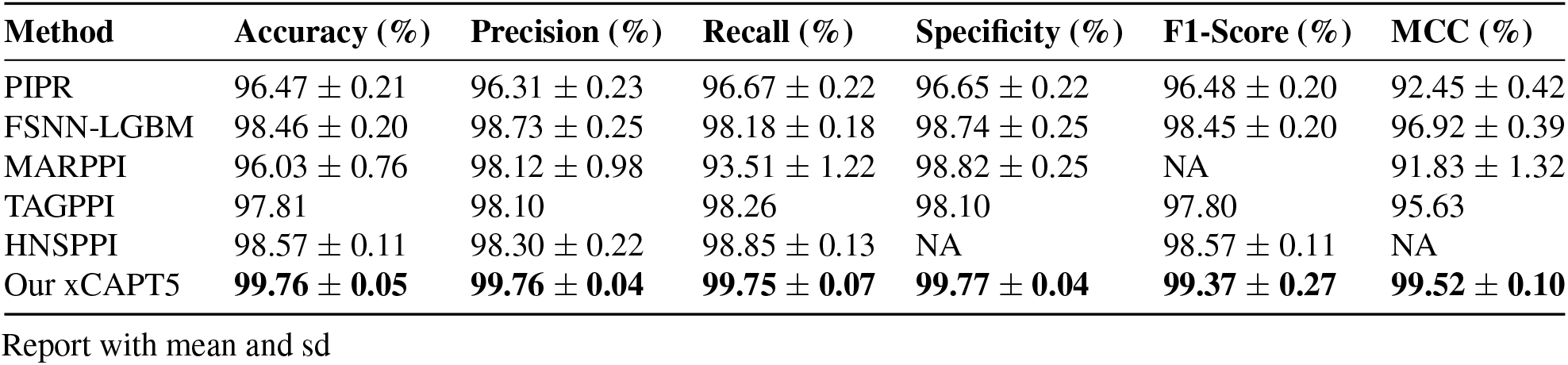
5-Fold cross-validation performances of methods on Guo dataset.

| Method | Accuracy (%) | Precision (%) | Recall (%) | Specificity (%) | F1-Score (%) | MCC (%) |
| --- | --- | --- | --- | --- | --- | --- |
| PIPR | 96.47 ± 0.21 | 96.31 ± 0.23 | 96.67 ± 0.22 | 96.65 ± 0.22 | 96.48 ± 0.20 | 92.45 ± 0.42 |
| FSNN-LGBM | 98.46 ± 0.20 | 98.73 ± 0.25 | 98.18 ± 0.18 | 98.74 ± 0.25 | 98.45 ± 0.20 | 96.92 ± 0.39 |
| MARPPI | 96.03 ± 0.76 | 98.12 ± 0.98 | 93.51 ± 1.22 | 98.82 ± 0.25 | NA | 91.83 ± 1.32 |
| TAGPPI | 97.81 | 98.10 | 98.26 | 98.10 | 97.80 | 95.63 |
| HNSPPI | 98.57 ± 0.11 | 98.30 ± 0.22 | 98.85 ± 0.13 | NA | 98.57 ± 0.11 | NA |
| Our xCAPT5 | **99.76 ± 0.05** | **99.76 ± 0.04** | **99.75 ± 0.07** | **99.77 ± 0.04** | **99.37 ± 0.27** | **99.52 ± 0.10** |

Report with mean and sd

Furthermore, on the Pan dataset, xCAPT5 significantly outperforms its closest competitors across all metrics, showcasing an accuracy of 99.77% with an exceptionally low standard deviation of 0.02%. The closest competitor, FSNN-LGBM, has a slightly lower accuracy of 99.50% but with a notably higher standard deviation of 0.28%, indicating less consistent results. The gap between xCAPT5 and its competitors is also significant. While FSNN-LGBM lags by a narrow margin of 0.27% in accuracy, this difference is amplified by the variation indicated by standard deviations. In precision, recall, and other metrics, xCAPT5 consistently ranks highest, almost always surpassing the 99.5% threshold with minimal variance.

### Generalized inference evaluation

We evaluate the generalization capacity of xCAPT5 and compared models by training them on human-centric data sets and subsequently testing on independent data sets. Our assessment encompasses a diverse range of test scenarios, spanning intra-species (human), cross-species (model organisms), and inter-species (human-virus) PPI datasets. The foundational training on human datasets equipped the models to discern patterns and features intrinsic to human protein interactions. By subjecting them to disparate test datasets, we aimed to ascertain the models’ proficiency in extrapolating their predictions beyond the confines of their training data. This rigorous analysis offers insights into the models’ competence in reliably predicting PPIs across varied biological contexts. Furthermore, it paves the way for the potential extrapolation of these models to species with scant or non-existent PPI data. In scenarios where specific PPI data is absent but protein sequence information is available, the models’ foundational training on human datasets can be harnessed to facilitate informed predictions.

#### Intra-species inference

The intra-species inference analysis presents the evaluation results of different methods on two distinct training datasets: the balanced training dataset Pan and the imbalanced training dataset Sledzieski. The performance of the methods is measured in terms of recall percentage on various test datasets.

Supplementary Table 1 shows the evaluation results for the intra-species dataset trained on the balanced Pan dataset. Across all test datasets (HPRD, DIP, HIPPIE HQ, HIPPIE LQ), xCAPT5 consistently achieves the highest recall. For instance, on the HPRD dataset, xCAPT5 achieves a recall of 96.16%, outperforming both PIPR (91.95%) and FSNN-LGBM (94.28%). The same trend is observed for other test datasets as well, with xCAPT5 consistently outperforming the other methods. xCAPT5 ranks first in terms of recall percentage for all of these datasets.

Supplementary Table 2 presents the evaluation results for the intra-species dataset trained on the imbalanced Sledzieski dataset. Despite the imbalance in both the training dataset and the test datasets, xCAPT5 again demonstrates superior performance. It achieves the highest recall on most test datasets. For example, on the DIP dataset, xCAPT5 achieves a recall of 67.64%, surpassing the recall of PIPR (30.79%) and FSNN-LGBM (48.71%). It achieves the highest recall on most test datasets, including HPRD, DIP, and HIPPIE HQ. However, it is worth noting that Topsy-Turvy achieves a slightly higher recall percentage of 51.22% on the HIPPIE LQ dataset compared to xCAPT5’s 40.92%.

#### Cross-species inference

The cross-species inference analysis shows the evaluation performance of different methods on cross-species datasets trained on two different training sets: Pan and Sledzieski. The test datasets represent various species: E. coli, Fly, Mouse, Worm, and Yeast.

In the Supplementary Table 3, where models are trained on the balanced training set Pan, we observe varying performance across the different methods and test datasets. D-SCRIPT consistently demonstrates the highest Precision, with values ranging from 70.64% (Yeast) to 85.47% (Mouse). It also achieves competitive F1-Scores, ranging from 33.88% (Yeast) to 53.68% (Fly), indicating a good balance between Precision and Recall. D-SCRIPT also performs well in terms of AUROC and AUPRC, achieving high values in most test datasets. Our model xCAPT5 shows the highest Recall values in several test datasets, such as Fly (83.08%) and Worm (71.02%). However, its Precision is relatively lower compared to D-SCRIPT.

In the Supplementary Table 4, where models are trained on the unbalanced training set Sledzieski, we can observe a decrease in overall performance compared to the first table. The Precision values of all methods are generally lower, indicating a higher number of false positives. However, xCAPT5 still shows the highest Precision, ranging from 9.18% (E. coli) to 9.45% (Yeast). Notably, the Recall values are consistently high across all methods and test datasets, ranging from 85.62% (Yeast) to 99.55% (E. coli) for xCAPT5.

#### Inter-species inference

In the Supplementary Table 5, the evaluation inference performance of our proposed model xCAPT5 and compared models on inter-species datasets trained on the balanced training set Pan is presented. The test datasets include Dengue, HIV, Hepatitis, Herpes, Influenza, Papilloma, SARS-CoV-2, and Zika.

Experimental results indicate that xCAPT5 generally performs the best across different test datasets. For example, in the Dengue test dataset, xCAPT5 achieves a precision of 9.21%, recall of 97.19%, F1-score of 16.83%, AUROC of 50.73%, and AUPRC of 9.44%. Our model demonstrates competitive performance across most test datasets. It achieves the highest Precision on the Hepatitis and Papilloma datasets and the highest Recall on the HIV dataset. Additionally, xCAPT5 achieves the highest F1-Score on the Zika dataset.

In the Supplementary Table 6, the evaluation inference performance of different methods on inter-species datasets trained on the unbalanced training set Sledzieski is presented. The test datasets are the same as in the Supplementary Table 5. Experimental results indicate that xCAPT5 performs well in most test datasets. For example, in the Dengue test dataset, xCAPT5 achieves a precision of 23.36%, recall of 35.66%, F1-score of 28.22%, AUROC of 54.90%, and AUPRC of 14.71%. Among the compared models, xCAPT5 consistently outperforms others in terms of Precision, Recall, and F1-Score on most test datasets. Notably, xCAPT5 achieves the highest Precision on the Hepatitis and Herpes datasets and the highest Recall on the HIV and Hepatitis datasets. It also obtains the highest F1-Score on the HIV and Influenza datasets.

### Stringent similarity evaluation

In this section, we assess the ability of our proposed model xCAPT5 to generalize to datasets with varying constraints on sequence similarity (Table 4). xCAPT5 stands out with its exceptional performance. It consistently achieves an accuracy of 99.72% and an F1 score of 99.61% across various sequence identities. This performance remains stable even when the sequence identity threshold tightens from 40% to just 1%. Such consistency indicates that xCAPT5 consistently delivers accuracy rates above 99.70% and F1 scores over 99.50%.

**Table 3.** 5-Fold cross-validation performances of methods on Pan dataset.

| Method | Accuracy (%) | Precision (%) | Recall (%) | Specificity (%) | F1-Score (%) | MCC (%) |
| --- | --- | --- | --- | --- | --- | --- |
| PIPR | 98.26 ± 0.02 | 98.68 ± 0.04 | 97.40 ± 0.04 | 97.93 ± 0.03 | 98.04 ± 0.02 | 96.49 ± 0.03 |
| FSNN-LGBM | 99.50 ± 0.28 | 98.48 ± 0.12 | 99.39 ± 0.54 | 99.58 ± 0.10 | 99.43 ± 0.32 | 98.98 ± 0.57 |
| Graph-BERT | 99.02 ± 0.13 | 98.94 ± 0.88 | 99.15 ± 0.95 | 98.57 ± 1.19 | 99.04 ± 0.10 | 98.00 ± 0.28 |
| Our xCAPT5 | **99.77 ± 0.02** | **99.75 ± 0.03** | **99.75 ± 0.02** | **99.80 ± 0.02** | **99.62 ± 0.06** | **99.55 ± 0.03** |

Report with mean and sd

**Table 4.** 5-Fold cross-validation performances of methods on stringent Chen multispecies datasets.

| Similarity | Methods | Accuracy (%) | F1-Score (%) |
| --- | --- | --- | --- |
| Any | PIPR | 98.19 | 98.17 |
|  | DeepTrio | 98.20 | 98.20 |
|  | TAGPPI | 99.15 | 99.15 |
|  | Our xCAPT5 | **99.72** | **99.61** |
| ≤ 40% | PIPR | 98.29 | 98.28 |
|  | DeepTrio | 97.83 | 97.98 |
|  | TAGPPI | 99.10 | 99.16 |
|  | Our xCAPT5 | **99.76** | **99.60** |
| ≤ 25% | PIPR | 97.91 | 98.08 |
|  | DeepTrio | 97.52 | 97.75 |
|  | TAGPPI | 98.99 | 99.06 |
|  | Our xCAPT5 | **99.74** | **99.61** |
| ≤ 10% | PIPR | 97.54 | 97.79 |
|  | DeepTrio | 97.32 | 97.62 |
|  | TAGPPI | 98.97 | 99.08 |
|  | Our xCAPT5 | **99.70** | **99.53** |
| ≤ 1% | PIPR | 97.51 | 97.80 |
|  | DeepTrio | 97.11 | 97.47 |
|  | TAGPPI | 98.89 | 98.89 |
|  | xCAPT5 | **99.73** | **99.60** |

Report with mean.

On the other hand, while PIPR, TAGPPI, and DeepTrio show commendable results, there’s a noticeable pattern: their performance metrics slightly decrease as the sequence identity requirements become stricter. This indicates that these models might face challenges when adapting to less familiar sequence spaces. The fluctuations in accuracy and F1-Score of xCAPT5 are minimal, with the most significant change being a mere 0.06% in accuracy. This consistent performance, even under tightening sequence similarity constraints, underscores xCAPT5’s robustness and superior generalization capabilities. Unlike many models that might falter under strict conditions, xCAPT5’s resilience is evident, suggesting that it’s adept at handling a broad spectrum of sequence identities without significant performance degradation.

### Hyperparameter effect

In this section, we assess the impact of hyperparameters on the performance of the xCAPT5 model, with a specific focus on the neural network architecture of xCAPT5. We employed a 5-fold cross-validation method on the Guo dataset to assess the neural architecture of xCAPT5 under different hyperparameter configurations. We note that increasing the number of kernel sizes from 2 to 3 leads to a significant performance improvement across multiple metrics. This suggests that a wider range of kernel sizes enables the model to detect a broader spectrum of patterns in the input data, enhancing overall performance. However, further increasing to four results in a decline in performance (Supplementary Figure 2). This deterioration can be attributed to increased complexity, making it harder for xCAPT5 to learn and generalize effectively. The model becomes more susceptible to capturing noise and irrelevant details, hindering its ability to discern relevant patterns and leading to decreased performance.

The depth of a Convolutional Neural Network (CNN), traditionally defined by the number of layers, plays a pivotal role in the model’s learning capacity. However, in the context of the xCAPT5 model, the depth is uniquely characterized by the number of modules, with each module representing a level of depth. The xCAPT5 model is composed of five such modules, signifying a depth of five. As the depth of the network increases, denoted by the number of modules in the xCAPT5 model (Supplementary Figure 1), there is a corresponding improvement in the model’s performance. The optimal performance is observed when the network comprises five modules. This optimal depth is influenced by certain parameters, such as the padding of the sequence length to 1200 and the use of a pooling size of 4.

Furthermore, our investigation encompasses the comparison of xCAPT5’s performance using different amino acid embeddings. In this regard, we discovered that leveraging the large protein language models like ProtT5-XL-U50, ProtT5-XL-BFD, ProtBert-BFD (12), and PlusRNN (21) provides superior results compared to traditional approaches like one-hot encoding and physicochemical concatenated with Skip-Gram embedding (Supplementary Figure 3). This highlights the importance of incorporating advanced protein language models in enhancing the predictive capabilities of xCAPT5.

## Conclusions

In this study, we proposed xCAPT5, a novel hybrid classifier that integrates the deep multi-kernel convolutional accumulated pooling siamese neural network (xCAPT5) with a XG-Boost model for improved protein-protein interaction (PPI) prediction. A defining aspect of xCAPT5’s architecture is its deployment of a deep multi-kernel convolutional neural network (CNN) within a Siamese structure. This approach allows the model to delve deeper into the intricate patterns encoded within and between protein sequences, enabling it to learn robust, discriminative features. The use of multiple kernels in the CNN allows the model to explore these patterns at different scales, increasing its ability to recognize a variety of sequence motifs, which can be critical for determining protein interactions.

Next, the integration of Global Average Pooling (GAP) and Global Max Pooling (GMP) further augments the model’s performance. Pooling operations reduce the spatial dimensions of the CNN output while retaining the most essential information. The use of both GAP and GMP in a concatenated manner ensures that the model retains the most representative features (as identified by GMP) while also preserving average trends in feature maps (as captured by GAP). This dual-pooling strategy enables the model to generate more comprehensive and meaningful representations of protein sequences. Finally, the incorporation of XGBoost, an optimized distributed gradient boosting library, as the second phase of xCAPT5 training, further boosts its performance. By learning on the representations generated by the neural network, XGBoost enhances the predictive capability of the model, leveraging its strength in handling mixed-type data, missing values, and its innate resistance to overfitting.

Our xCAPT5 model exhibits state-of-the-art performance as demonstrated through rigorous cross-validation on multiple well-established datasets. In the cross-validation analysis, xCAPT5 was evaluated alongside PIPR, FSNN-LGBM, TAGPPI, HNSPPI, Graph-BERT, and MARPPI on three datasets: Martin, Guo, and Pan. xCAPT5 consistently outperformed the other methods across all datasets, achieving the highest on each metrics.

The experimental results of the cross-species inference analysis revealed interesting insights. When trained on the balanced dataset Pan, xCAPT5 consistently demonstrated strong performance across multiple evaluation metrics, including Precision, Recall, F1-Score, AUROC, and AUPRC. This suggests that xCAPT5 is capable of effectively predicting protein-protein interactions across different species when provided with a balanced training dataset.

The inter-species inference analysis further highlights the state-of-the-art performance of xCAPT5 trained on Pan and Sledzieski. xCAPT5 consistently outperformed the compared SOTA models in terms of Precision, Recall, and F1 on most test datasets, regardless of the training dataset. Notably, xCAPT5 achieved the highest Precision on the Hepatitis and Papilloma datasets and the highest Recall on the HIV and Hepatitis datasets. These findings emphasize the effectiveness of xCAPT5 in predicting protein-protein interactions across different species, regardless of the training data.

Finally, the experimental results from the stringent similarity evaluation demonstrate the excellent generalization ability of xCAPT5 in comparision with several SOTA models, including PIPR, TAGPPI, and DeepTrio. Interestingly, xCAPT5 can consistently achieve the highest accuracy and F1 among all compared SOTA models across different sequence similarity thresholds. This is a valuable characteristic of xCAPT5 as it allows reliable PPI predictions even in cases where sequence homology is low.

In future work, we aim to explore the interpretability of xCAPT5 predictions through contact maps, a binary, two-dimensional representation that illustrates the contacts or interactions between two different proteins. It visualizes all pairwise spatial contacts between amino acid residues from the two proteins. These maps represent the interactions between distinct protein complexes, may hold the key to interpreting xCAPT5’s predictive power. Additionally, by incorporating attention mechanisms, we could discern which aspects of the contact maps are most critical for accurate PPI prediction. Furthermore, the development of a multichannel input scheme to accommodate diverse information sources such as evolutionary profiles, physicochemical properties, or secondary structure predictions can also enhance the model. Collectively, these explorations could enhance our understanding of protein-protein interactions and the predictive power of xCAPT5.

## ACKNOWLEDGEMENTS

We would like to thank Miss Mai-Anh Hang Vo at VNU University of Science for her contributions to scientific illustration.

## Supplementary data on hyperparameter effects

**Supplementary Fig. 1.**
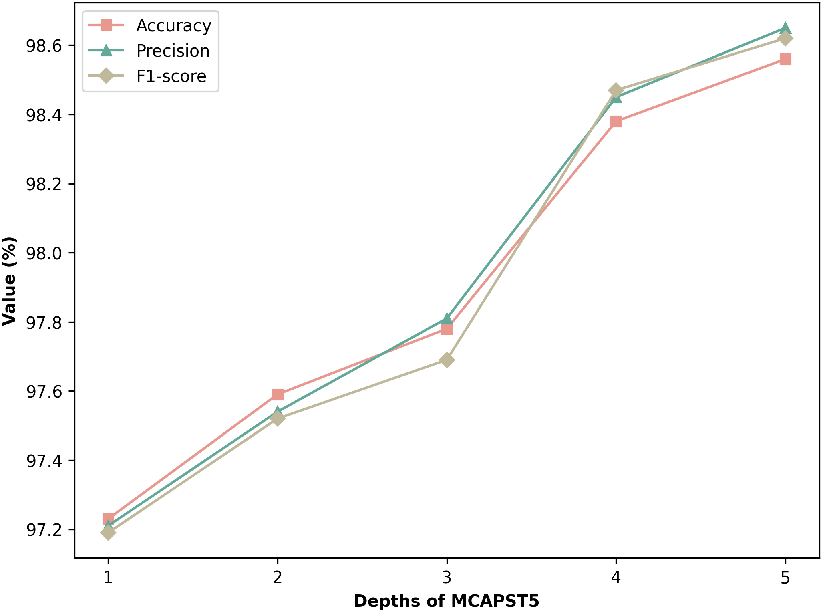
Evaluation hyperparameter depth of xCAPT5.

**Supplementary Fig. 2.**
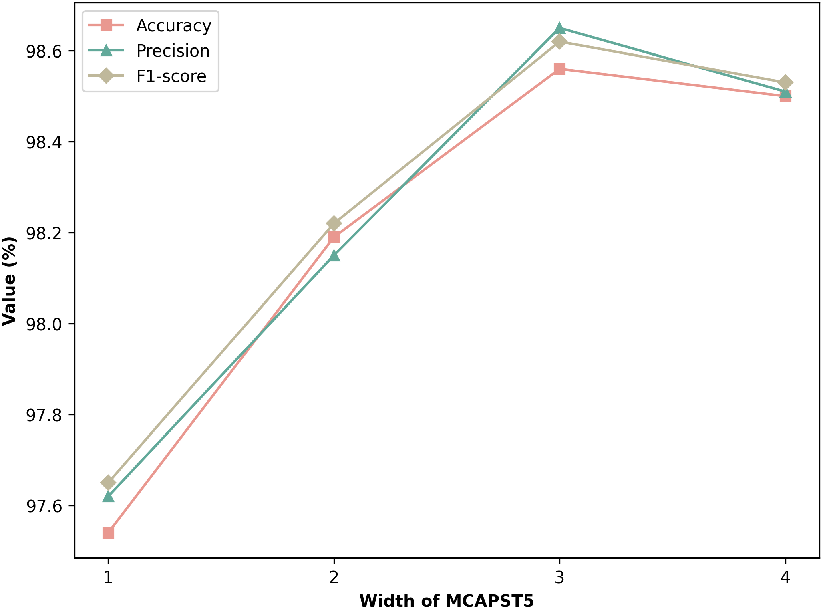
Evaluation hyperparameter width of xCAPT5.

**Supplementary Fig. 3.**
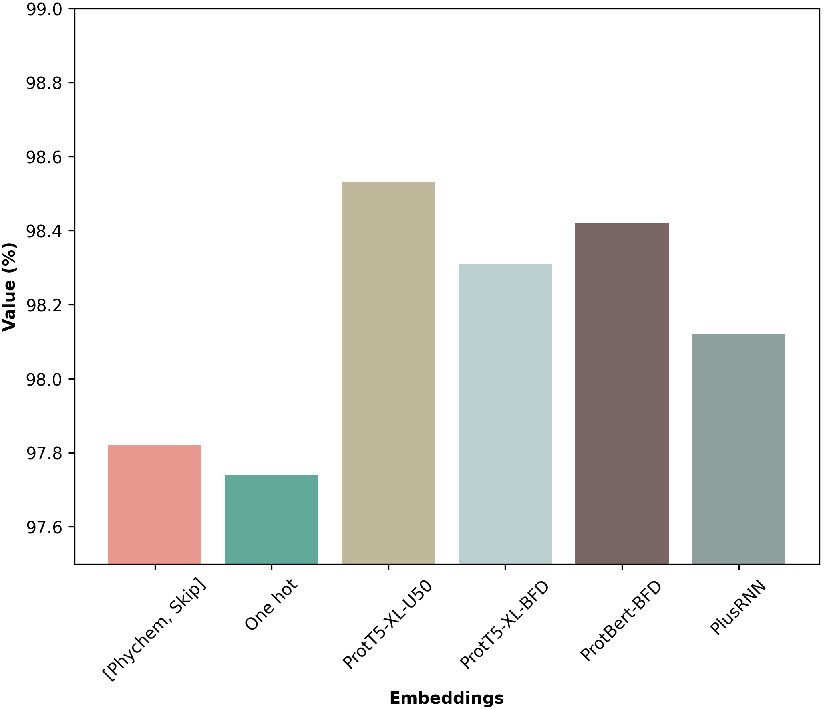
Evaluation hyperparameter embeddings of xCAPT5.

## Supplementary data on intra-species inference

**Supplementary Table. 1.**
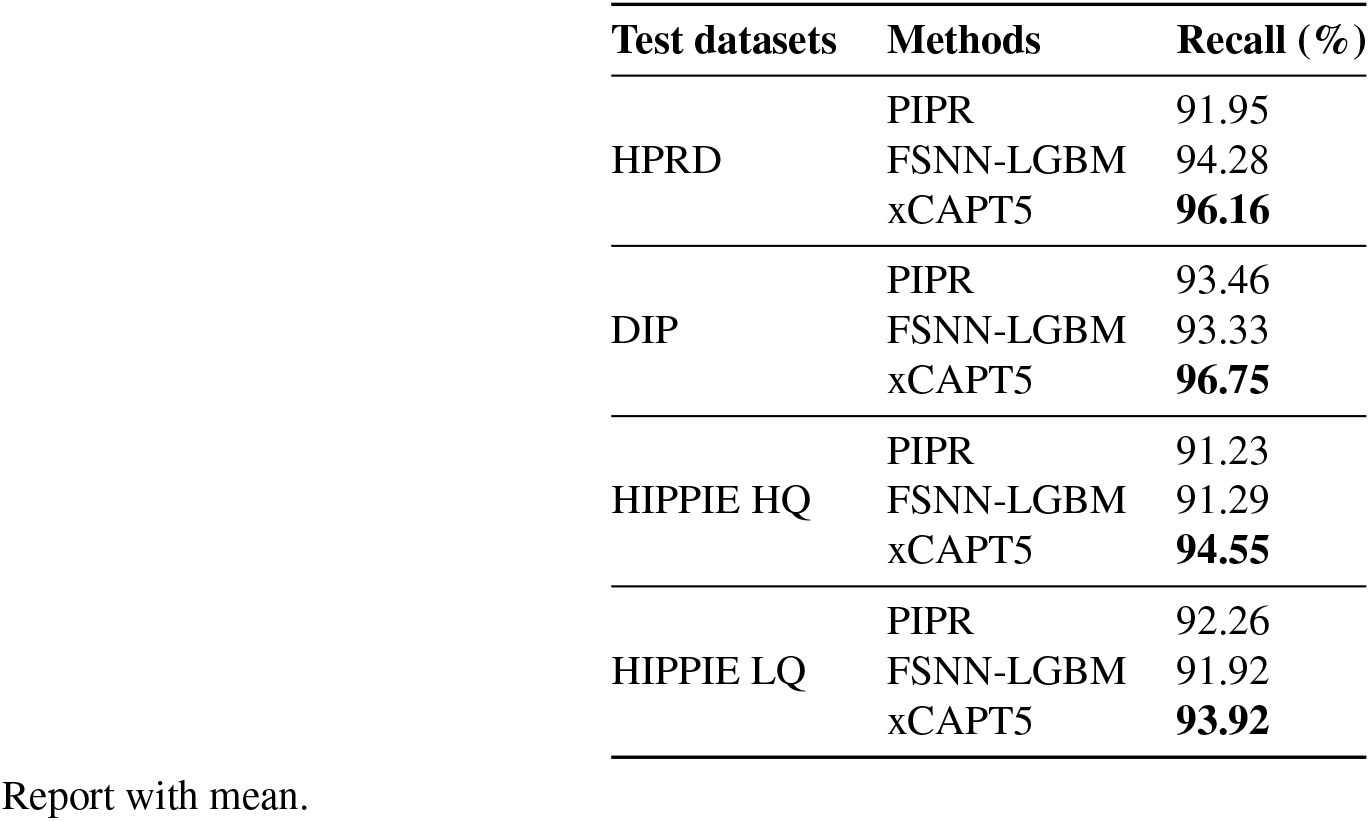
Evaluation inference performance of methods on intra-species dataset trained on Pan dataset.

| Test datasets | Methods | Recall (%) |
| --- | --- | --- |
| HPRD | PIPR | 91.95 |
|  | FSNN-LGBM | 94.28 |
|  | xCAPT5 | **96.16** |
| DIP | PIPR | 93.46 |
|  | FSNN-LGBM | 93.33 |
|  | xCAPT5 | **96.75** |
| HIPPIE HQ | PIPR | 91.23 |
|  | FSNN-LGBM | 91.29 |
|  | xCAPT5 | **94.55** |
| HIPPIE LQ | PIPR | 92.26 |
|  | FSNN-LGBM | 91.92 |
|  | xCAPT5 | **93.92** |

Report with mean.

**Supplementary Table. 2.** Evaluation inference performance of methods on intra-species dataset trained on Sledzieski dataset.

| Test datasets | Methods | Recall (%) |
| --- | --- | --- |
| HPRD | PIPR | 22.84 |
|  | FSNN-LGBM | 40.93 |
|  | D-SCRIPT | 12.62 |
|  | Topsy-Turvy | 51.22 |
|  | xCAPT5 | **58.53** |
| DIP | PIPR | 30.79 |
|  | FSNN-LGBM | 48.71 |
|  | D-SCRIPT | 11.44 |
|  | Topsy-Turvy | 56.67 |
|  | xCAPT5 | **67.64** |
| HIPPIE HQ | PIPR | 32.24 |
|  | FSNN-LGBM | 46.41 |
|  | D-SCRIPT | 12.54 |
|  | Topsy-Turvy | 45.19 |
|  | xCAPT5 | **58.84** |
| HIPPIE LQ | PIPR | 22.83 |
|  | FSNN-LGBM | 38.54 |
|  | D-SCRIPT | 7.23 |
|  | Topsy-Turvy | **51.22** |
|  | xCAPT5 | 40.92 |

Report with mean.

## Supplementary data on cross-species inference

**Supplementary Table. 3.** Evaluation inference performance of methods on cross-species dataset trained on Pan dataset.

| Test datasets | Methods | Precision (%) | Recall (%) | F1-Score (%) | AUROC (%) | AUPRC (%) |
| --- | --- | --- | --- | --- | --- | --- |
| E. Coli | PIPR | 8.91 | 90.21 | 16.22 | 46.88 | 8.53 |
|  | FSNN-LGBM | 8.67 | 92.14 | 15.84 | 47.51 | 8.73 |
|  | xCAPT5 | **9.18** | **99.55** | **16.79** | **50.47** | **9.17** |
| Fly | PIPR | 8.59 | 87.70 | 15.65 | 44.39 | 8.13 |
|  | FSNN-LGBM | 8.74 | 92.59 | 15.98 | 48.01 | 8.78 |
|  | xCAPT5 | **9.29** | **96.70** | **16.95** | **51.13** | **9.71** |
| Mouse | PIPR | 8.72 | 89.76 | 15.89 | 46.78 | 8.53 |
|  | FSNN-LGBM | 8.85 | 93.12 | 16.16 | 48.59 | 8.86 |
|  | xCAPT5 | **9.36** | **97.46** | **17.07** | **51.52** | **9.35** |
| Worm | PIPR | 8.61 | 90.24 | 15.73 | 43.35 | 7.98 |
|  | FSNN-LGBM | 8.81 | 93.63 | 16.11 | 48.36 | 8.83 |
|  | xCAPT5 | **9.31** | **96.50** | **16.98** | **51.24** | **9.30** |
| Yeast | PIPR | 8.52 | 85.62 | 15.50 | 43.83 | 8.03 |
|  | FSNN-LGBM | 8.86 | 93.29 | 16.19 | 48.68 | 8.88 |
|  | xCAPT5 | **9.45** | **94.78** | **17.18** | **51.98** | **9.43** |

Report with mean.

**Supplementary Table. 4.** Evaluation inference performance of methods on cross-species dataset trained on Sliedzieski dataset.

| Test datasets | Methods | Precision (%) | Recall (%) | F1-Score (%) | AUROC (%) | AUPRC (%) |
| --- | --- | --- | --- | --- | --- | --- |
| E. Coli | PIPR | 55.81 | 17.30 | 26.41 | 70.85 | 31.59 |
|  | FSNN-LGBM | 19.10 | 24.95 | 21.63 | 57.19 | 11.59 |
|  | D-SCRIPT | **75.54** | 38.30 | 50.83 | **86.03** | **53.54** |
|  | Topsy-Turvy | 39.81 | 46.40 | 42.85 | 64.30 | 37.22 |
|  | xCAPT5 | 44.41 | **64.95** | **52.75** | 78.41 | 32.03 |
| Fly | PIPR | 52.17 | 14.44 | 22.62 | 73.21 | 29.56 |
|  | FSNN-LGBM | 22.62 | 30.42 | 25.95 | 60.01 | 13.21 |
|  | D-SCRIPT | **81.56** | 40.00 | 53.68 | 84.17 | 59.69 |
|  | Topsy-Turvy | 63.01 | 75.62 | 68.74 | 90.07 | **67.66** |
|  | xCAPT5 | 62.77 | **83.08** | **71.51** | **91.08** | 53.69 |
| Mouse | PIPR | 67.06 | 34.94 | 45.94 | 85.08 | 52.53 |
|  | FSNN-LGBM | 32.05 | 51.56 | 39.53 | 70.32 | 20.93 |
|  | D-SCRIPT | **85.47** | 34.48 | 49.14 | 79.92 | 54.91 |
|  | Topsy-Turvy | 59.03 | 71.04 | 64.48 | 86.41 | **60.80** |
|  | xCAPT5 | 53.14 | **84.78** | **65.33** | **87.64** | 54.35 |
| Worm | PIPR | 62.81 | 15.64 | 25.04 | 76.54 | 34.80 |
|  | FSNN-LGBM | 25.49 | 29.50 | 27.35 | 60.44 | 13.93 |
|  | D-SCRIPT | **85.03** | 32.98 | 47.53 | 80.21 | 56.31 |
|  | Topsy-Turvy | 70.55 | 63.85 | 67.04 | 85.20 | **62.98** |
|  | xCAPT5 | 77.52 | **71.02** | **74.13** | **86.48** | 59.69 |
| Yeast | PIPR | 37.71 | 10.56 | 16.50 | 71.30 | 22.68 |
|  | FSNN-LGBM | 22.27 | 24.76 | 23.45 | 58.06 | 12.36 |
|  | D-SCRIPT | **70.64** | 22.28 | 33.88 | 78.89 | 40.46 |
|  | Topsy-Turvy | 48.32 | 54.36 | 51.16 | 76.20 | 43.14 |
|  | xCAPT5 | 62.87 | **58.82** | **60.78** | **79.67** | **44.72** |

Report with mean.

## Supplementary data on inter-species inference

**Supplementary Table. 5.**
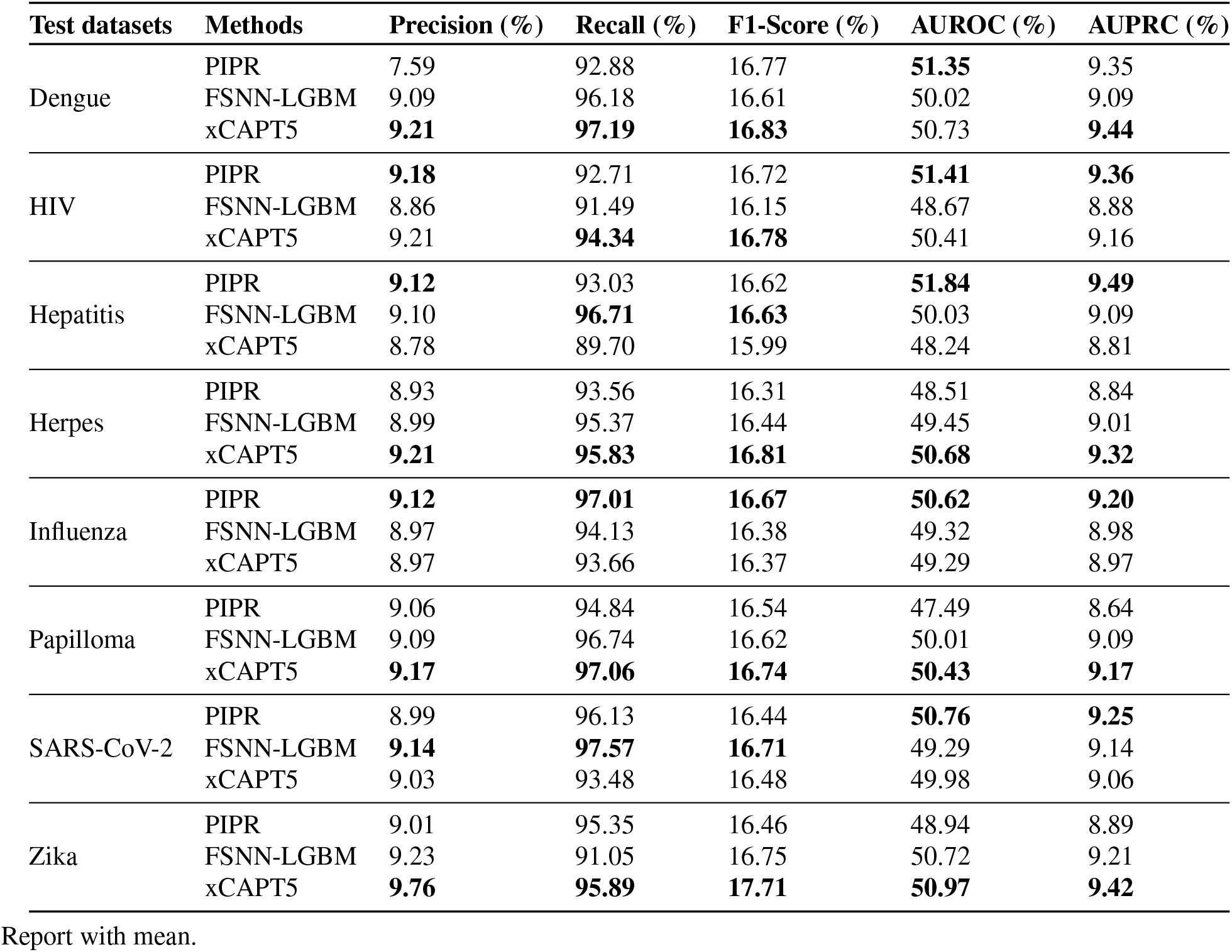
Evaluation inference performance of methods on inter-species dataset trained on Pan dataset.

**Supplementary Table. 6.** Evaluation inference performance of methods on inter-species dataset trained on Sledzieski dataset.

| Test datasets | Methods | Precision (%) | Recall (%) | F1-Score (%) | AUROC (%) | AUPRC (%) |
| --- | --- | --- | --- | --- | --- | --- |
| Dengue | PIPR | 24.68 | 6.87 | 10.62 | **62.31** | 15.32 |
|  | FSNN-LGBM | 18.31 | 33.82 | 24.02 | 59.67 | 12.33 |
|  | D-SCRIPT | 17.98 | 1.45 | 2.68 | 61.25 | 13.16 |
|  | Topsy-Turvy | 22.01 | 22.71 | 22.35 | 61.08 | **16.14** |
|  | xCAPt5 | **23.36** | **35.66** | **28.22** | 54.90 | 14.71 |
| HIV | PIPR | 24.25 | 9.10 | 13.19 | 63.91 | 16.02 |
|  | FSNN-LGBM | 17.57 | 32.58 | 22.83 | 58.64 | 11.86 |
|  | D-SCRIPT | 19.46 | 4.63 | 7.47 | 50.11 | 10.96 |
|  | Topsy-Turvy | 16.87 | 7.19 | 10.08 | 53.44 | 11.04 |
|  | xCAPt5 | **27.14** | **36.10** | **30.98** | **64.20** | **17.61** |
| Hepatitis | PIPR | 20.84 | 1.96 | 3.58 | 51.42 | 11.03 |
|  | FSNN-LGBM | 15.94 | 21.46 | 18.09 | 54.94 | 10.49 |
|  | D-SCRIPT | **32.68** | 0.97 | 1.89 | **59.43** | 12.92 |
|  | Topsy-Turvy | 20.61 | 7.87 | 11.39 | 50.99 | 10.24 |
|  | xCAPt5 | 20.71 | **22.46** | **21.55** | 56.93 | **13.70** |
| Herpes | PIPR | 22.88 | 6.21 | 9.74 | 56.89 | **12.94** |
|  | FSNN-LGBM | 16.06 | **26.87** | 20.10 | 56.41 | 10.97 |
|  | D-SCRIPT | **27.54** | 0.90 | 1.75 | **58.96** | 12.74 |
|  | Topsy-Turvy | 18.51 | 8.39 | 11.53 | 50.36 | 10.55 |
|  | xCAPt5 | 22.23 | 21.79 | **22.01** | 57.08 | 12.35 |
| Influenza | PIPR | 21.62 | 6.16 | 9.43 | 62.54 | 14.42 |
|  | FSNN-LGBM | 18.63 | **34.65** | 24.22 | 59.75 | 12.39 |
|  | D-SCRIPT | 14.06 | 1.19 | 2.19 | 56.17 | 11.49 |
|  | Topsy-Turvy | 19.82 | 13.01 | 15.66 | 55.91 | 11.55 |
|  | xCAPt5 | **22.67** | 30.80 | **26.12** | **63.17** | **16.61** |
| Papilloma | PIPR | 16.62 | 2.92 | 4.94 | 55.61 | 11.37 |
|  | FSNN-LGBM | 16.43 | **26.33** | 20.23 | 56.47 | 11.02 |
|  | D-SCRIPT | **22.87** | 1.47 | 0.92 | 52.13 | 10.67 |
|  | Topsy-Turvy | 15.95 | 8.58 | 11.14 | 51.60 | 10.35 |
|  | xCAPt5 | 20.12 | 24.12 | **21.93** | **58.32** | **13.04** |
| SARS-CoV-2 | PIPR | 13.11 | 3.58 | 5.56 | 55.92 | 11.13 |
|  | FSNN-LGBM | 15.47 | **27.41** | 19.77 | 56.23 | 10.84 |
|  | D-SCRIPT | **21.48** | 1.21 | 2.33 | 59.02 | 13.08 |
|  | Topsy-Turvy | 15.79 | 7.52 | 10.19 | 55.56 | 10.59 |
|  | xCAPt5 | 17.90 | 22.89 | **20.09** | **60.29** | **14.81** |
| Zika | PIPR | 13.59 | 6.96 | 8.77 | 57.21 | 12.57 |
|  | FSNN-LGBM | 15.56 | **30.84** | **20.67** | 57.09 | 11.15 |
|  | D-SCRIPT | **42.47** | 7.27 | 12.40 | **69.06** | **20.57** |
|  | Topsy-Turvy | 14.72 | 19.40 | 16.72 | 57.03 | 12.17 |
|  | xCAPt5 | 16.59 | 25.39 | 20.07 | 51.90 | 13.56 |

Report with mean.

